# Retrosplenial cortex is necessary for spatial and non-spatial latent learning in mice

**DOI:** 10.1101/2021.07.21.453258

**Authors:** Ana Carolina Bottura de Barros, Liad J. Baruchin, Marios C. Panayi, Nils Nyberg, Veronika Samborska, Mitchell T. Mealing, Thomas Akam, Jeehyun Kwag, David M. Bannerman, Michael M. Kohl

## Abstract

Latent learning occurs when associations are formed between stimuli in the absence of explicit reinforcement. Traditionally, latent learning in rodents has been associated with the creation internal models of space. However, increasing evidence points to roles of internal models also in non-spatial decision making. Whether the same brain structures and processes support the creation of spatially-anchored or non-spatial internal models via latent learning, is an open question. To address this question, we developed a novel operant box task that allows to test spatial and non-spatial versions of a flavour-based sensory preconditioning paradigm. We probed the role of the retrosplenial cortex, a brain area associated with spatial cognition and subjective value representation, in this task using precise, closed-loop optogenetic silencing during different task phases. We show that the retrosplenial cortex is necessary for both spatial and non-spatial latent learning in mice. We further demonstrate that the requirement of retrosplenial cortex is limited to the preconditioning phase of the task. Our results provide insight into the specific role of the retrosplenial cortex in latent learning, demonstrate that latent learning plays a general part in the creation of internal models, independent of spatial anchors, and provide a novel avenue for studying model-based decision making.

## Introduction

Decision making refers to the process of selecting a particular option among a set of alternatives expected to produce different outcomes ^1^. One way the brain is thought to aid decision making is through updating an internal model ^2,3^. Such updates may be produced by latent learning, in which information about the causal relationships between stimuli is stored in the absence of appetitive/aversive outcomes. While latent learning was initially thought to help create internal maps of space ^4^ there is a growing appreciation that it may be also important for creating more general, non-spatial cognitive maps ^5,6^. Whether different brain regions specialise in supporting spatial or non-spatial forms of latent learning remains an open question.

Sensory preconditioning (SPC) can be used to assess latent learning experimentally where two neutral stimuli, A and B, are presented together in a preexposure phase, typically followed by conditioning stimulus B to an appetitive or aversive outcome, before testing the animal’s response to the never-conditioned stimulus A ^7^. At test, subjects respond to A as if it predicted the outcome, even though they were never explicitly paired. This suggests that latent learning about the associative relationship between A-B resulted in an internal model that, when combined with learning that B -> Outcome, allows for an inference that A-B -> Outcome.

Several brain structures have been implicated in rodent SPC, including hippocampus ^8,9^, perirhinal ^10,11^, entorhinal ^5,12^ orbitofrontal ^13,14^ and retrosplenial cortex (RSC) ^15–17^. The RSC has mostly been associated with spatial cognition ^18–22^ but may also be involved in processing non-spatial information ^23–25^. This made the RSC an ideal place to test whether latent learning is a broadly employed process that it is used in updating not just spatially anchored models but more fundamentally also models lacking spatial markers. Furthermore, in contrast to previous work ^15–17^, our task design allowed precise control of foreground and background cues as well as removing possible spatial confounds by not using auditory or visual cues thus allowing us to test spatial and non-spatial versions of the same SPC task while precisely silencing RSC using closed-loop optogenetics.

## Results

A viral construct containing the red-shifted cruxhalorhodopsin, Jaws ^26^ under control of the CaMKII⍺ RSC in adult male wild-type mice (Figure 1A). This gave robust expression of Jaws-GFP throughout the dorsal RSC (Supplementary Figure 1). We used 32-contact linear silicon probe recordings in anaesthetised mice to confirm that neurons in the agranular and granular part of the RSC could be silenced by red light delivered for varying durations (150 to 1000 ms) corresponding to those in behavioural tasks (Figure 1B). We compared single unit recordings at different depths of the RSC between groups of Jaws-injected experimental mice and control mice that received equivalent injections of a virus lacking Jaws (AAV5-CamKIIα-eYFP). We found that activation of Jaws produced temporally precise and reversible silencing of RSC neurons (Figure 1C-H; three-way ANOVA (light × depth × group); interaction of light and group: *F*(1,18) = 11.348, *p* = 0.003, main effect of group: *F*(1,18) = 17.614, *p* = 0.001; *n* = 2000 trials from one experimental and one control mouse) and that the efficacy of the optogenetic silencing did not vary with recording depth (three-way ANOVA; all depth interactions *F*(1,2) < 2.003, *p* > 0.150). After an initial rebound, activity was not significantly different to controls after around 150 ms.

**Figure 1:**
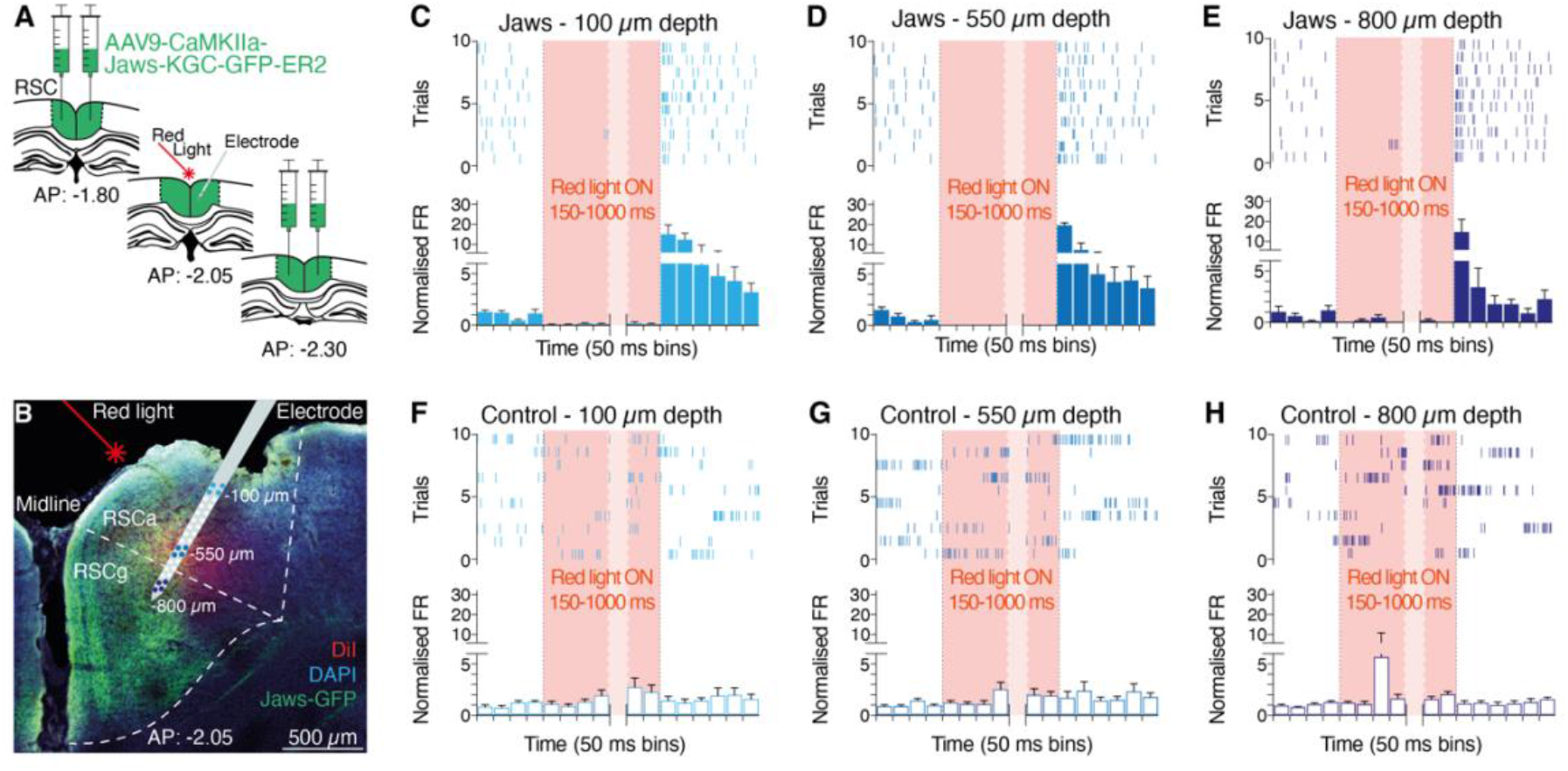
Optogenetics enables temporally precise and reversible silencing of retrosplenial cortex (RSC) *in vivo*. **(A)** Adeno-associated virus containing the Jaws-KGC-GFP-ER2 (“Jaws”) or e-YFP (“Control”) construct under the control of the CaMKII promotor was used. Virus was injected into two sites per hemisphere of the RSC (target spread indicated in green; black dotted lines: RSC boundaries). Virus injection resulted in Jaws-GFP or e-YFP expression in the dorsal RSC. **(B)** A 32-contact silicon electrode was used to record single unit activity before, during and after illumination with red light delivered via an optical fibre. Image shows a fixed section of a brain after recording (Jaws-GFP in green; DAPI in blue; approximate location of electrode based on DiI track in red is indicated; electrode width is not to scale; coloured circles on probe represent the different recording depths shown in panels C-H; white dotted line: boundaries of granular (RSCg) and agranular (RSCa) parts of RSC). **(C-E)** Illumination of RSC neurons in Jaws-expressing mice resulted in a reversible reduction in spontaneous spiking frequency in RSCa and RSCg. **(F-H)** Illumination of RSC neurons in GFP-expressing control mice had no effect on spontaneous spiking frequency. Variable duration light ON epoch indicated in red. Representative spike raster for ten neurons at each recording depth as well as normalised mean firing rate for each 50 ms bin are shown. Error bars represent *SEM*.

For behavioural testing, a lightweight red LED ^27^ was implanted between the four virus injection sites to illuminate RSC (Figure 2A). We verified LED implant placement and the level of viral spread through histological analyses after behavioural testing and confirmed that there were no significant differences in LED placement and viral spread (Supplementary Figure 1).

**Figure 2:**
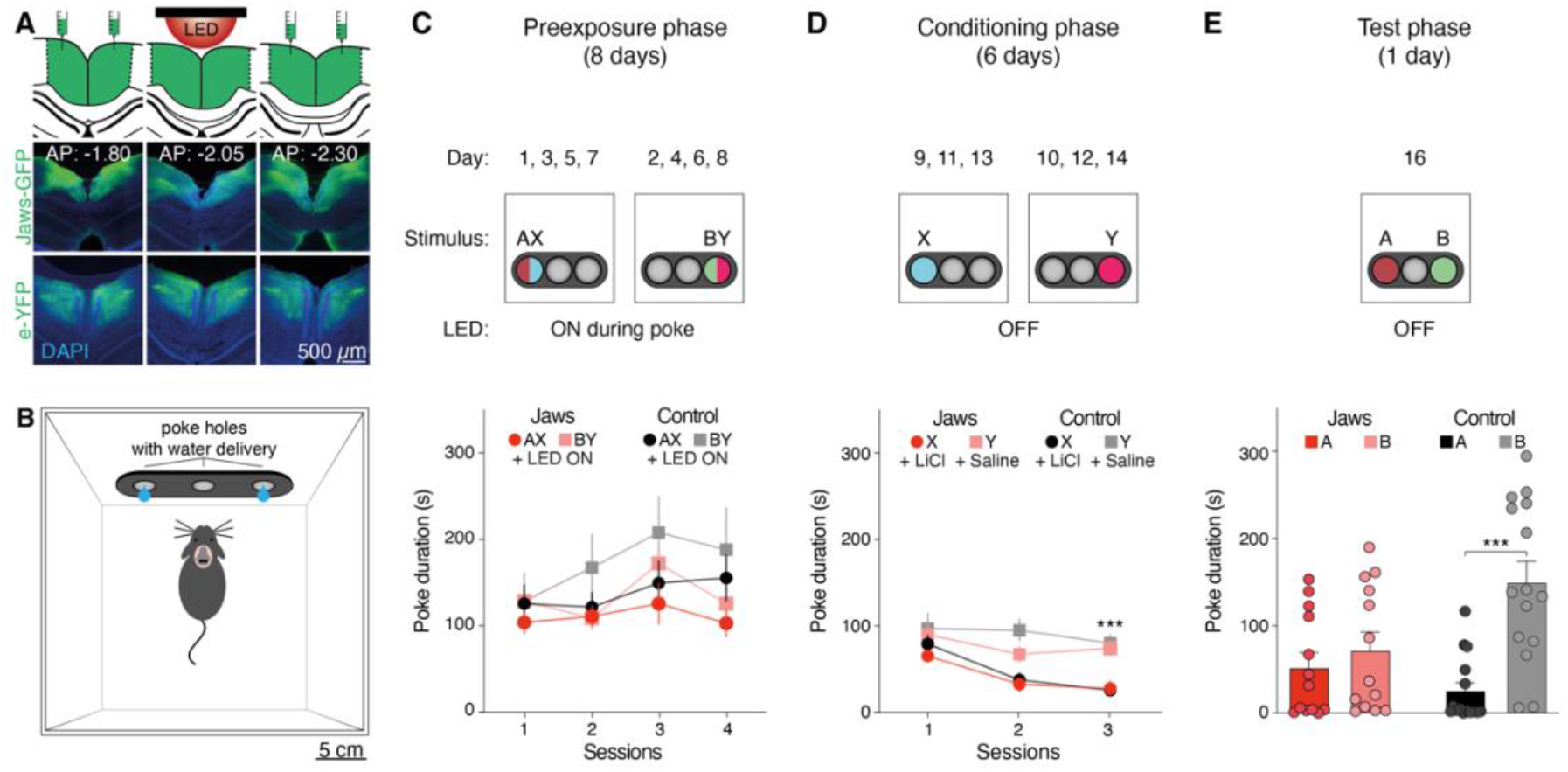
Latent learning of neutral stimulus-stimulus pairs requires retrosplenial cortex. **(A)** Virus injection into two sites per hemisphere resulted in Jaws-GFP or e-YFP expression in the entire dorsal retrosplenial cortex. Approximate location of the two injection sites per hemisphere and LED implant are indicated (not to scale). Expression at approximate location of implant is indicated. **(B)** Mice were trained on a modified sensory preconditioning task in an operant box using the left and right nose-poke ports, each able to dispense small amounts of flavoured water. Duration of pokes was used to estimate liquid consumption. **(C)** Upper panel: During the preexposure phase, mice received a mixture of two flavours on alternate days from different nose-poke ports over the course of eight days. Mice therefore experienced each flavour combination four times, shown as Session. Flavours indicated as dark red (A), blue (X) green (B) and pink (Y) circles in nose-poke ports. Infrared sensors in the nose-poke ports enabled closed-loop activation of the implanted red LED every time the mouse poked to consume flavoured liquid. Lower panel: There was no difference in the water consumption between Jaws and e-YFP-control animals over the course of the preexposure phase. **(D)** Upper panel: During the conditioning phase, the LED was OFF, and mice received only one part of the mixture of two flavours on alternate days from different nose-poke ports over the course of six days. After a session of drinking flavour X, mice received a lithium chloride (LiCl) injection *i.p.* After a session of drinking flavour Y, mice received a saline injection *i.p.* Lower panel: Consumption of flavour X significantly reduced over the course of the conditioning phase. **(E)** Upper panel: During the test phase, the LED was OFF, and mice could freely choose to receive either flavour A or B from different nose-poke ports. Lower panel: There was no difference in consumption of flavour A or B in Jaws-injected animals but a significant rejection for flavour A, which was previously paired with the LiCl-conditioned flavour X, in e-YFP-control mice. Total poke duration is shown. Error bars represent *SEM*. *** *p* ≤ 0.001, two-way ANOVA.

To study the role of RSC in the latent learning of spatial or non-spatial stimulus-stimulus associations, we adapted a flavour-based sensory preconditioning (SPC) task ^28^ for use in an operant box set-up. Jaws and control groups of mice could obtain small amounts of differently flavoured water from nose-poke ports that were fitted with infrared beam-breaks. This enabled us to activate the implanted LED in a closed-loop fashion with millisecond precision and to record an animal’s decision for one or the other flavour as the duration of time spent in a nose-poke port (Figure 2B). The overall task structure consisted of four phases: pretraining, preexposure, conditioning and test (Figure 2C-E, top panels). In the spatial version of the task, different flavour stimuli were delivered from spatially separate nose-poke ports. In the non-spatial version, flavours were always delivered from the same, central nose-poke port until the test phase.

In the pretraining phase animals were placed into the operant box for up to 30 minutes while receiving unflavoured water from the nose-poke ports. This accustomed them to the background cues (e.g. operant box, house lights, enclosure fan noise) and the experimental procedure (e.g. loading animals into the boxes, nose-poke triggered water delivery system).

During the preexposure phase of the spatial version of the task, mice were for the first time exposed to the foreground cues consisting of flavoured water. A combination of flavours A and X (AX) were offered from one side nose-poke port on one day and a combination of flavours B and Y (BY) from the opposite side nose-poke port on the next day. This was repeated over the course of eight days. The implanted LED was turned on only for a pseudo random duration (150–1000 ms) while AX or BY were consumed in the preexposure phase. Both groups (*n_Control_* = 15; *n_Jaws_* = 13) showed equivalent levels of water consumption for all flavour combinations indicating that there was no flavour preference or effect of Jaws activation on drinking behaviour during preexposure (Figure 2C). During the conditioning phase, mice were offered water flavoured with only one component of the combinations experienced during the preexposure phase: flavour X from the nose-poke port where AX was offered during preexposure and flavour Y from the nose-poke port where BY was offered during preexposure. Immediately after removing mice from the operant boxes, those presented with flavour X received an intraperitoneal injection of LiCl, which triggers short episodes of diarrhoea that become associated with the recently experienced flavour. Flavour Y exposure was followed with a saline injection. This was repeated over the course of six days. Both Jaws and Control groups successfully acquired conditioned flavour aversion to X (three-way ANOVA, flavour by session interaction: *F*(2,48) = 7.848, *p* = 0.001; Figure 2D). There was no difference in the behavioural measures for Jaws and Control mice throughout preexposure and conditioning phases (see Detailed Statistics *in Supplementary Materials*). On the subsequent test day, we evaluated the ability of mice to infer a reduced value of flavour A based on the previous experience of the AX flavour combination followed by the *devaluation* of flavour X. We offered mice a choice between water with flavour A or flavour B during one final session from the nose-poke ports corresponding to where AX and BY had been delivered during the preexposure phase. Control animals displayed a strong preference for flavour B over A, while Jaws animals showed no preference for flavour (Figure 2E). We observed a significant interaction between flavour and group (*F*(1,24) = 4.788, *p* = 0.039), a main effect of flavour (*F*(1,24) = 16.301, *p* < 0.001) and a significant difference in consumption of B over A for Controls (*F*(1,24) = 21.445, *p* < 0.001) but not for Jaws (*F*(1,24) = 1.56, *p* = 0.224). There was no difference in the total amount of water consumed between groups. These results demonstrate that Jaws mice were unable to infer a lower value of flavour A compared to B as a result of the earlier silencing of RSC during the experience of combinations of flavours.

The non-spatial version of the task repeated the experiment with a modification: during the preexposure and conditioning phases, flavoured water was always only offered from the central nose-poke port, eliminating potential spatial foreground cues. As before, during the preexposure phase, both groups (*n_Control_* = 14; *n_Jaws_* = 16) showed equivalent levels of flavoured water consumption for all flavour combinations (Figure 3B). During the conditioning phase, both Jaws and Control groups successfully acquired conditioned flavour aversion to X (three-way ANOVA, main effects of session and of flavour (Flavour: *F*(2,52) = 16.524, *p* < 0.001; Session: *F*(2,52) = 26.984, *p* < 0.001; Figure 3C). Again, there was no difference between Jaws and Control mice during preexposure and conditioning phases (see Detailed Statistics *in Supplementary Materials*). On the test day Control mice displayed a strong preference for flavour B over A, while the Jaws group did not show a preference for either flavour (Figure 3D). We observed a significant interaction between flavour and group (*F*(1,26) = 4.949, *p* = 0.035) and a main effect of flavour (*F*(1,26) = 7.098, *p* = 0.013). Simple effects revealed that Control group displayed a significant difference in consumption of B over A (*F*(1,26) = 11.204, *p* = 0.002), but Jaws did not (*F*(1,26) = 0.104, *p* = 0.750).

**Figure 3:**
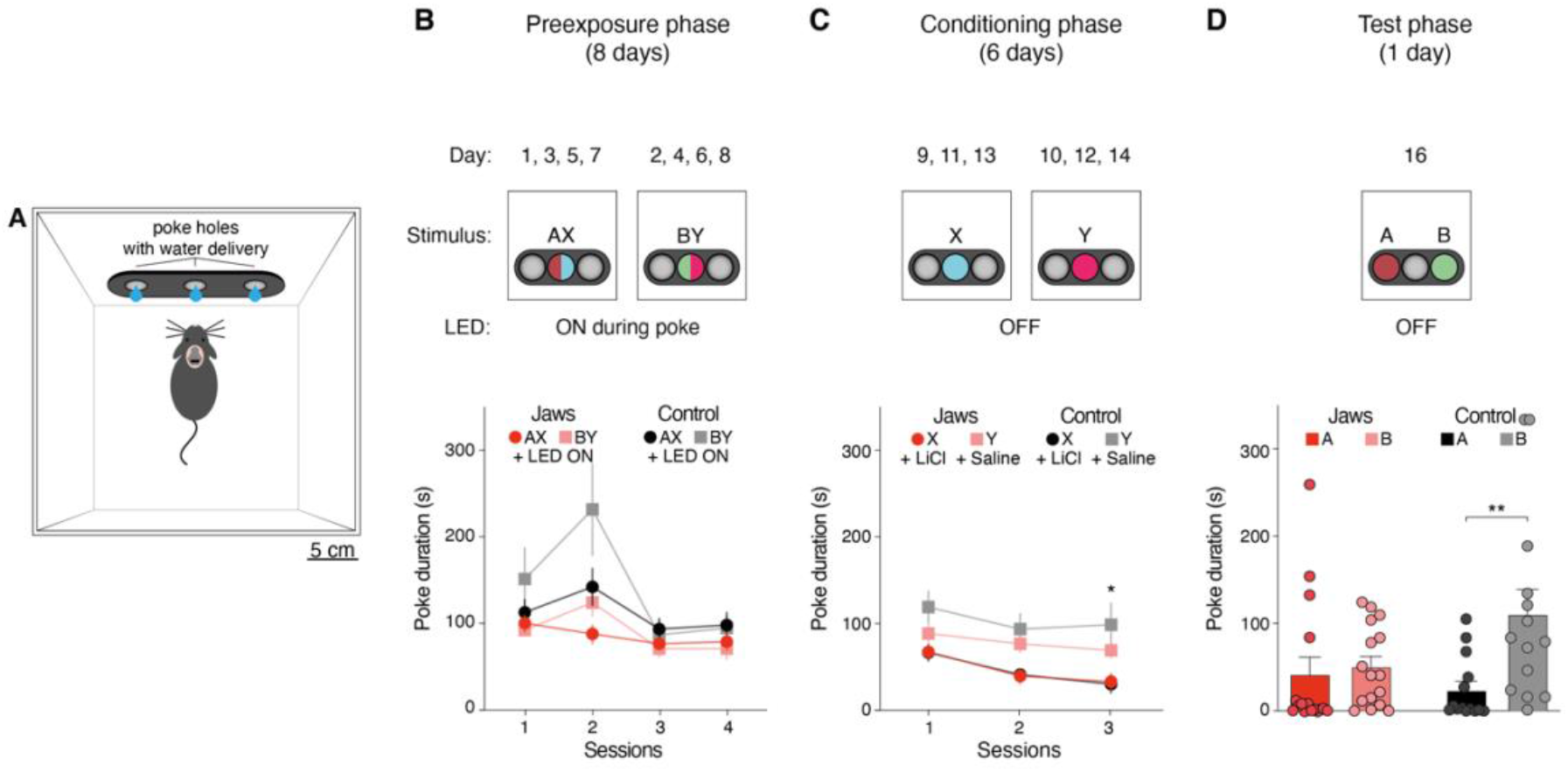
Retrosplenial cortex-dependent latent learning of stimulus-stimulus pairs without spatial landmarks. **(A)** In the non-spatial version of the task, mice were trained in a modified operant box using only the central nose-poke port during the preexposure and conditioning phases and the left and right nose-poke ports during decision making in the test phase. **(B)** Upper panel: During the preexposure phase, mice received a mixture of two flavours on alternate days from the same, central nose-poke port over the course of eight days. Mice therefore experienced each flavour combination four times, shown as Session. Flavours indicated as dark red (A), blue (X) green (B) and pink (Y) circles in nose-poke ports. Infrared sensors in the nose-poke ports enabled closed-loop activation of the implanted red LED every time the mouse poked to consume flavoured water. Lower panel: There was no difference in the liquid consumption between Jaws and e-YFP-control animals over the course of the preexposure phase. **(C)** Upper panel: During the conditioning phase, the LED was OFF, and mice received only one part of the mixture of two flavours on alternate days from the same, central nose-poke port over the course of six days. After a session of drinking flavour X, mice received a lithium chloride (LiCl) injection *i.p.* After a session of drinking flavour Y, mice received a saline injection *i.p.* Lower panel: Consumption of flavour X significantly reduced over the course of the conditioning phase. **(D)** Upper panel: During the test phase, the LED was OFF, and mice could freely choose to receive either flavour A or B from different side nose-poke ports. Lower panel: There was no difference in consumption of A or B flavoured water in Jaws-injected animals but a significant rejection for flavour A, which was previously paired with the LiCl-conditioned flavour X, in e-YFP-control mice. Total poke duration is shown. Error bars represent *SEM*. * *p* ≤ 0.02, ** *p* = 0.002, two-way ANOVA.

## Discussion

Here we show that the retrosplenial cortex makes a temporally specific and critical contribution to latent learning. This critical contribution was not limited to spatial processing.

Previous work demonstrating impaired SPC to visual and auditory cues following long-term inhibition of RSC via electrolytic lesions and chemogenetic manipulations ^15–17^ was unable to dissociate the roles of background (e.g. operant box, house lights, enclosure fan noise) and foreground (e.g. nose-poke ports, flavoured water) cues. This was because RSC activity was disrupted throughout the entire training session. We addressed this issue by allowing the animals to experience the background cues in a pretraining phase prior to exposure of the foreground cues in the preexposure phase and through the use of closed-loop optogenetics that limited the disruption of RSC activity to the duration of foreground cue exposure. Using flavoured stimuli instead of visual or auditory cues, we were able to include a choice test that increased test sensitivity.

Chronic recordings ^29–31^ and long-term RSC lesions in rodents and humans ^21,32,33^ all point to a role of the RSC in the representation of spatial information. However, we find that RSC is also required for latent learning in the non-spatial version of the task which is unlikely to be solved in a model-free manner. This suggests a fundamental role of RSC in model-based decision making. This notion is supported by recent results showing that RSC encodes inferred value-related signals more stably than primary sensory or association cortices ^23^. Together, these findings additionally point to latent learning supporting the generation of internal models beyond spatial markers. Our work shows RSC is required for latent learning alongside other brain areas and offers a novel, more sensitive approach to study the latent learning processes supporting model-based decision making.

## Methods

### Subjects

All animal husbandry and experimental procedures were approved and conducted in accordance with the United Kingdom Animals (Scientific Procedures) Act 1986 under project and personal licenses from the Home Office. Four C57Bl/6 male mice were used in electrophysiological experiments. For behavioural experiments, a total of 64 C57Bl/6 male mice, 8–16 weeks old, ~30 g (Biomedical Services, Oxford, UK) received injections, implants and training. Mice were housed in groups in a climate-controlled vivarium (lights on 7:00 to 19:00). The holding room temperature was 23±1 degree Celsius and humidity was set to 40±10%. The experiments were conducted during the light portion of the photoperiod. Mice were fed low-fat mouse chow in the form of pellets and had *ad libitum* access to food but access to water was restricted during the experiment. Animals were kept at 85–90% of free-feeding weight for the duration of the behavioural experiments. All behavioural experiments and histology were performed with the experimenter blind to the identity of the mice.

### Surgery

Under anaesthesia, six to seven weeks old mice received surgery for viral injection for opsin expression and a light-emitting diode (LED) implantation for optogenetic stimulation. Anaesthesia was induced via inhalation of 4% isoflurane (Zoetis, Leatherhead, UK) at 1 L/min. When mice were fully anesthetized, they were placed in a stereotaxic frame (Kopf instruments, Tujunga, CA). The depth of anaesthesia was monitored by checking pedal withdrawal reflex and level of Isoflurane was kept at 0.5–1% at 0.4 L/min during surgery. Before surgery, mice received injections of meloxicam (5 mg/kg, Metacam, Boehringer Ingelheim International GmbH, Ingelheim am Rhein, Germany) and vetergesic (0.1 mg/kg, Ceva Animal Health Ltd, Amersham, UK). They also received a marcaine (AstraZeneca, Cambridge, UK) injection under the scalp.

A circular incision was made into the scalp, the skull was cleaned, and the periosteum removed in preparation for the craniotomy. A 1.8 mm diameter craniotomy centred on the midline and at –2.05 mm from Bregma was performed leaving the dura intact. Injections of either rAAV9/CamKII-Jaws-KGC-GFP-ER2 (4.8×10^12^ vg/ml) or rAAV5/CamKII-EYFP (7.4×10^12^ vg/ml) (University of North Carolina Vector Core, Chapel Hill, NC) were made using an automated injector (Nanoject II, Drummond Scientific, Broomall, PA) with in-house made, bevelled micropipettes fashioned from thin-walled borosilicate glass (3.5 inches, Drummond Scientific). Mice received injections at two sites per hemisphere (ML: ± 0.3 mm; AP: –1.8 and –2.3 mm). At each site, 50 nL injections were made at different depths (from dura: –0.8/–0.6/–0.4/–0.2 mm) at 50 nL/min. The injection needle was left in place for 4 min before being removed from each site.

Finally, custom made red (630 nm wavelength) LED headstages (Newbury Electronics Ltd, Newbury, UK) were cemented (Simplex Cement, Kemdent, Swindon, UK; Superbond C&B, Sunmedical, Moriyama, Japan) on top of the craniotomy. After surgeries, animals were kept in a thermoregulated chamber until they recovered from anaesthesia. They were then transferred to post-operative cages.

### Electrophysiology recordings

Injections of rAAV9/CaMKII-Jaws-KGC-GFP-ER2 were carried out as described above. After at least three weeks of expression had passed, electrophysiological recordings were performed.

Anaesthesia was induced using Urethane (10% dissolved in saline, Sigma Aldrich, Gillingham, UK) at 1–1.5 mg/kg body weight. Once depth of anaesthesia was ascertained by the lack of tail-pinch and leg-withdrawal reflexes, mice were placed in a stereotaxic frame. In the injected mice, a craniotomy was made on top of RSC region using a dental drill. In some cases, mice were used for recordings after being tested behaviourally. In these cases, prior to the craniotomy, after ensuring a suitable depth of anaesthesia the head-cap was removed by drilling carefully around it, melting the cement with acetone and dampening the implant with saline and then pulling it carefully off the skull.

Then, a 32-channel single shank silicon probe (A1×32-Poly2-5mm-50s-177; Neuronexus, Ann Arbor, MI) was inserted into the right hemisphere at AP: –2.05 mm, ML: –0.30 mm at approximately 45-degree angle to the cortical surface. The probe was slowly lowered to about DV: –800 μm and recordings started after 10 minutes wait. Recordings were made with light delivered from a laser via a multi-mode fibre optic cable (200 μm diameter, 0.39 NA; Thorlabs, Ely, UK). To match the light intensity of the LED implant (50 mW) and account for possible losses in power, for example, because of bone regrowth, we pooled recordings using three light intensities (50, 20, or 5 mW at the tip of a fibre optic cable). The fibre optic cable was held just above the pial surface to mimic illumination with the implanted LED These recordings used the same temporal stimulation regime as the one used for behavioural tests (pseudorandom, 150–1000 ms).

### Apparatus

Behavioural experiments were conducted in eight identical custom-built operant boxes (12 cm × 12 cm × 12 cm) made of Perspex and placed into sound attenuating chambers (SAC) with an exhaust fan. The operant boxes featured a roofed ceiling with an opening through which a ~15 cm long light-weight cable attached to a rotary commutator could be attached and connected to the head-mounted LED. The back wall of the operant box featured three nose-poke ports with backlight, infrared beam detection and a solenoid valve-controlled liquid delivery spout. House lights were mounted on the side walls of the SAC. All boxes were controlled through open-source behavioural hardware and, PyControl ^34^ (OEPS Electrónica e Produção, Alges, Portugal). The flavoured stimuli used were 10% sucrose solution (A or B; Sigma Aldrich), 0.15 M saline solution (A or B; Sigma Aldrich), 0.01 M hydrochloric acid (X or Y; Fisher Scientific, Loughborough, UK), and 60.0 μM quinine (X or Y; Sigma Aldrich). When the liquid solutions were presented in combination, they were formulated to retain these concentrations.

### Behavioural Procedure

#### Phase 1, Pretraining

Water restricted mice had the implanted LEDs connected and were placed into the operant box for 30 min or until they had consumed 30 water rewards delivered from one of the three nose-poke ports. Subjects had to poke to get a 15 μL delivery of drinking water. There was a timeout of 2 s before the next reward was delivered if the animal was still in the nose-poke port or tried to poke again. This pattern of reward delivery was repeated in all following phases. In the Spatial Sensory Preconditioning only the side nose-poke ports were used, therefore pretraining consisted of two sessions, one for each side nose-poke port, delivered on one day. In the Non-Spatial version of the task, all three nose-poke ports were used, therefore three pretraining sessions were completed. On the first day of pretraining animals received two sessions one for each of the side nose-poke ports. The following day a single session with delivery from the centre nose-poke port.

#### Phase 2, Preexposure

The second phase was composed of eight days. Each morning, mice had 30 min exposure to flavour compounds AX or BY on alternate days. For the Spatial Sensory Preconditioning, AX would be delivered through one of the side nose-poke ports and BY, the next day, on the opposite side nose-poke port. Side of presentation of solutions was counterbalanced between animals, but constant over days. For the Non-Spatial version of the task, both were delivered through the centre nose-poke port on alternate days. Nose-poke port backlights were lit up to indicate that reward was available. Subjects had to poke to get a 15 μL delivery of compound. Poking triggered the reward after a pseudorandom interval (10–35 ms) and the optogenetic stimulation by turning on the implanted LED (at 50 mW). Light would be ON for a pseudorandom duration (150–1000 ms). Rewards were delivered every 2000 ms if the animal remained in the nose-poke port. Every time a reward was delivered, stimulation would be triggered as described above.

Each afternoon, approximately four hours after training, mice were given access to water for 30 minutes.

#### Phase 3, Conditioning

During phase 3, for six days, mice were allowed to drink from the centre nose-poke port X only or Y only on alternate days in 30-minute sessions. When the sessions finished, after drinking X, all mice received an injection of 0.15 M LiCl (40ml/kg, Sigma Aldrich). After drinking Y, mice received a 0.9% saline injection. For the Spatial version X and Y were delivered from the side nose-poke ports, respectively where AX and BY had been delivered. For the Non-Spatial SPC, both X and Y were delivered from the centre nose-poke port on alternate days.

#### Phase 4, Test

For the test session, A and B were presented simultaneously for 30 min. Mice had a choice of drinking either from the left or from the right nose-poke ports. A and B location was counterbalanced for the Non-Spatial version of the task. For the Spatial SPC, A and B followed the presentation side for AX and BY.

For all phases, consumption was measured by calculating the duration of nose pokes. Counterbalancing factors (stimulus identity, side of presentation) were included in the statistical analysis when any interaction was observed.

### Histology

Animals were anaesthetised with sodium pentobarbital intraperitoneally (60 mg/kg, Pentoject, AnimalCare, York, UK) and transcardially perfused initially with 0.1 M phosphate buffer saline (PBS, Sigma Aldrich) followed by 4% paraformaldehyde (PFA, TAAB Laboratories, Aldermaston, UK) in 0.1 M phosphate buffer (pH 7.4). Brains were kept overnight in PFA at 4 degree Celsius and then changed to 0.1 M phosphate buffer saline (PBS) and stored at 4 degree Celsius until histological procedures.

Perfused brains were cut into 50 μm slices using a microtome (Leica Biosystems, Milton Keynes, UK) and one in every six slices was chosen for analysis and imaging. Brain slices were incubated in 4’,6-diamidino-2-phenylindole (DAPI, Acros Organics–Thermo Fisher Scientific, Geel, Belgium) and mounted on microscope slides in mounting medium (Vectashield, Vector Labs, Burlingame, CA) and covered for imaging.

Images of whole brain slices for analysis of virus expression were obtained from a multichannel fluorescence microscope (EVOS FL Auto Imaging System, Thermo Fisher Scientific, Waltham, MA) with an automatic objective at 10X magnification. Detailed images of slices from brains used for electrophysiological recordings were obtained from a laser scanning confocal microscope (FV1000, Olympus, Hamburg, Germany), equipped with an air 20X/0.75 NA and an oil-immersion 40X/0.8 NA objective.

### Data Analysis

For the electrophysiology analysis, multiunit activity was detected by filtering the signal with a 500 Hz highpass Butterworth filter. Any threshold crossing higher than 5 SD of baseline was considered as a spike. Activity was collected in 10 sessions of 100 trials each. PSTH was calculated by averaging sessions split into 50 ms bins. The change in neural activity was calculated by normalising the activity to baseline, equivalent to the 200 ms pre-light stimulation. Analysis of variance was used to compare activity sessions for effect of light ON (spiking activity during 200 ms pre-light stimulation and during the first 200 ms after the light came on) × recording depth (100 μm, 550 μm and 800 μm below pial surface) × experimental group (Jaws and Control) using GLM Repeated Measures in SPSS Statistics (v. 25.0.0.1, IBM Corporation, Cambridge, UK). The same model was used to compare within subjects repeated measures factors when the analysis was expanded to include more animals.

Nose poke data was collected throughout the experiments. Number of pokes and duration were calculated. Custom written Python codes were used for analysis of operant boxes behavioural output. For both preconditioning experiments, analysis of behaviour with within subjects repeated measures factors were analysed using GLM Repeated Measures in SPSS Statistics. Initially, counterbalancing factors (stimulus identity) were added as a covariate in the model but removed i f they did not interact with Group.

Image analysis software ImageJ (Fiji, https://fiji.sc) was used to analyse and quantify viral expression areas in histologically processed brain slices. Viral expression area was manually determined and traced with ImageJ for area calculation. Infected areas outside of the RSC region were also traced and calculated. An effort was made to keep the observer blind to the experimental condition, but the viral constructs used lead to a different expression pattern. The opsin Jaws is expressed in the membrane due to trafficking sequences KGC and ER2 while in the controls e-YFP expression is cytoplasmatic. Allen Brain Atlas (http://mouse.brain-map.org/static/atlas) coronal schemes were used to estimate expression per RSC area in ImageJ.

## Acknowledgements

This work was supported by a Medical Research Council Grant (MR/T004363/1) to M.M.K. and a grant of the Korea Health Technology R&D Project through the Korea Health Industry Development Institute (KHIDI), funded by the Ministry of Health & Welfare, Republic of Korea (HI19C0646) to J.K.A.C.B.B. was supported by a CNPq - Brazil studentship. M.M.K. was supported by an Oxford Early Career Research Fellowship. Tai-Ying Lee assisted with some of the conditioning experiments. Viral vectors were obtained from Dr. R. Jude Samulski and the University of North Carolina Vector Core.

## Author contributions

Study design: A.C.B.B., D.M.B., M.M.K., M.C.P., T.A.; data acquisition: A.C.B.B., L.J.B., M.M.K., N.N., V.S.; analysis: A.C.B.B., L.J.B., M.T.M., M.M.K., M.C.P.; manuscript writing: A.C.B.B., L.J.B., M.M.K.; manuscript editing: A.C.B.B., M.C.P., T.A, L.J.B., J.K., D.M.B., V.S., M.M.K.; resources: D.M.B., J.K., M.M.K.

## Competing interests

The authors have no competing interests to report.

## Materials & Correspondence

Correspondence and material requests should be addressed to A.C.B.B. or M.M.K.

## Supplementary Information

### Detailed Statistics

#### Experiment 1: preexposure and conditioning stimuli from side nose-poke ports

##### Preexposure

There was an interaction between group, session and flavour (*F*(3,60) = 2.982, *p* = 0.038). This was due to a transient preference for BY in the Control group shown by analysis of simple effects of the interaction (Control, Day 3: *F*(1,20) = 4.969, *p* = 0.037). Given that the identity of the flavours used in the liquid compounds was fully counterbalanced, this does not appear to be a flavour preference. It was likely due to mice being run at a slightly later time which increased thirst on that day. Importantly, there were no differences for flavours on days 1, 2, or 4 (Control: All *F* < 4.969, *p* > 0.083; Jaws: All *F* < 3.175, *p* > 0.09). Furthermore, a two-way ANOVA on data from session 4 revealed no interaction between group and flavour (*F*(1,20) = 0.044, *p* = 0.836) indicating that by the last session there was no difference in both group’s drinking behaviour.

##### Conditioning

There was no significant main effect of group (*F*(1,24) = 0.887, *p* = 0.356) or any interaction with group (All *F* < 0.827, *p* > 0.444), suggesting a robust acquisition of taste aversion to X by both groups. Simple effects of favour by session interaction: session 1: *F*(1,24) = 4.400, *p* = 0.047; session 2: *F*(1,24) = 40.981, *p* < 0.001; session 3: *F*(1,24) = 38984, *p* < 0.001. No significant differences were observed on session 1 for the analysis of the flavour X session interaction (Control: *F*(1,24) = 1.539, *p* = 0.227; Jaws: *F*(1,24) = 2.927, *p* = 0.1).

##### Test

The overall total amount consumed by both groups did not differ as demonstrated by the absence of a main effect of group (*F*(1,24) = 1.877, *p* = 0.183). A main effect of flavour: *F*(1,24) = 16.301, *p* < 0.001) was observed.

#### Experiment 2: preexposure and conditioning stimuli from central nose-poke port

##### Preexposure

Statistical analysis revealed no interaction between group, session and flavour (*F*(3,66) = 2.061, *p* = 0.114), but there was an interaction between flavour and session, between session and group and main effects of session and of flavour (All *F* > 4.245, *p* < 0.046). Because we observed these effects, an analysis of simple effects of the three-way interaction was run and showed that Control animals transiently preferred BY over AX on the first 2 sessions (Control - Session 1 and 2: *F* > 5.281, *p* < 0.031 and Jaws - Session 1 and 2 *F* < 2.306, *p* > 0.143). Importantly, no preference was observed on the last two sessions (Control and Jaws - Session 3 and 4: *F* < .318, *p* > 0.579). This was likely due to experimental error as some animals were still adapting to water regulation. In particular, ANOVA of flavour and group variables on the last day showed no interaction (*F*(1,22) = 0.086, *p* = 0.772) and no main effects (Flavour: *F*(1,22) = 0.224, *p* = 0.641; Group: *F*(1,22) = 2.954, *p* = 0.1).

##### Conditioning

There was no flavour by session interaction; all sessions: *F*(2,52) = 0.804, *p* = 0.453; session 1: *F*(1,26) = 9.921, *p* = 0.004; session 2: *F*(1,26) = 11.455, *p* = 0.002; session 3: *F*(1,26) = 14.090 *p* = 0.001; There were no interactions with group and no main effect of group (Interactions: *F* < 1.105, *p* > 0.339; Main effect: *F*(1,26) = 0.042, *p* = 0.839). Thus, highlighting that both in preexposure and conditioning phases there were no group differences.

##### Test

A main effect of pairing was also present (*F*(1,26) = 7.098, *p* = 0.013). Again, the absence of a main effect of group (*F*(1,24) = 1.877, *p* = 0.183) indicates that the overall total amount drank by both groups did not differ.

### Histology

As our electrophysiological experiments confirmed that RSC activity is disrupted by Jaws activation, we set out to test the effect of RSC optogenetic disruption on sensory associations. 28 C57Bl6 mice received injections of either a viral vector for expression of Jaws or a control vector and had a red LED implanted on top of the skull (Figure 2A). Supplementary Figure 1 shows expression spread area in representative animals covering the full range from minimum area expression to maximum one. In summary, between AP coordinates –0.8 to –3.8 mm from Bregma, control animals expressed the control virus in 36.0% ± 3.3% of the RSC area and Jaws expression was about 36.6% ± 3.4% (one way ANOVA, main effect of AP level: *F*(1,5) = 7.258; *p* = 0.044). We found no difference between experimental groups (one way ANOVA, AP level by Group interaction: *F*(1,5) = 0.315; *p* = 0.599; main effect of Group: *F*(1,5) = 0.665; *p* = 0.452). If we consider the RSC area immediately below the craniotomy where the LED was implanted the levels of Jaws expression increase to 46.1% ± 4.3%.

---

**Supplementary Figure 1:**
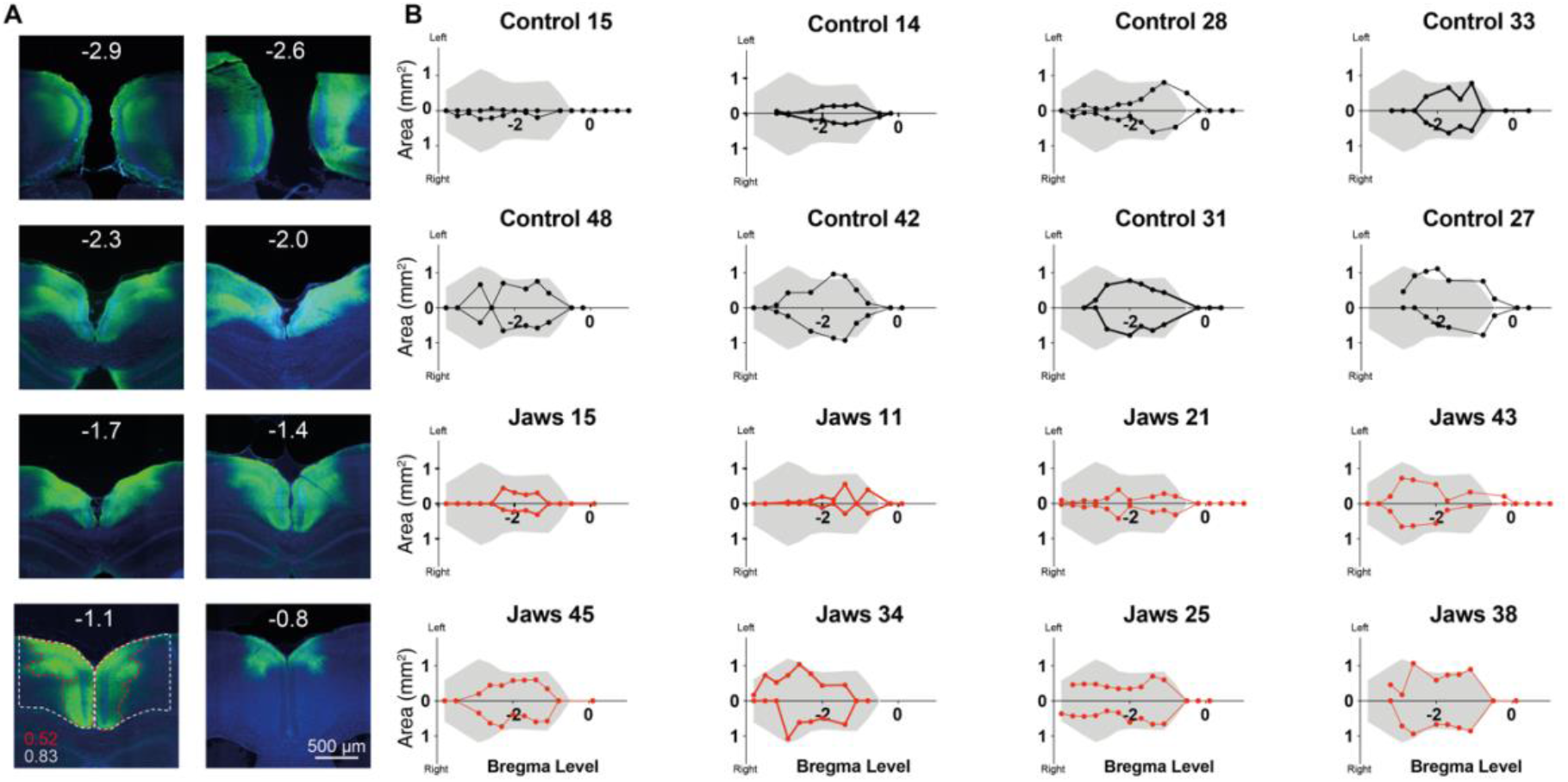
Robust viral expression throughout dorsal RSC. **(A)** Example of viral expression spread in a mouse injected with the Jaws viral construct (mouse ID: Jaws #35). AP levels are shown in white at the top. AP –1.1 illustrates how viral spread was estimated by comparing measurements for areas of fluorescence (red dotted lines) with the area of the RSC as defined by the Allen Brain Atlas (white dotted lines) for 14 coronal sections each separated by 300 μm. Corresponding area values in mm^2^ are shown. (**B**) Representative examples illustrating minimum (top left) to maximum (bottom right) expression levels in eight control and eight Jaws mice. The shaded grey area represents the projection of the RSC area to the dorsal, cortical surface, and plotted along the posterior-anterior axis from approximately lambda (−4 mm) to bregma (0 mm).

## Notes

### Competing Interest Statement

The authors have declared no competing interest.

## References

1. Lee, D. Decision making: from neuroscience to psychiatry. Neuron 78, 233–48 (2013).

2. Doll, B. B., Simon, D. A. & Daw, N. D. The ubiquity of model-based reinforcement learning. Curr Opin Neurobiol 22, 1075–81 (2012).

3. Sutton, R. S. & Barto, A. G. Reinforcement Learning: An Introduction. Ieee T Neural Networ 9, 1054–1054 (1998).

4. Tolman, E. C. Cognitive maps in rats and men. Psychol Rev 55, 189 (1948).

5. Stouffer, E. M. & Klein, J. E. Lesions of the lateral entorhinal cortex disrupt non-spatial latent learning but spare spatial latent learning in the rat (Rattus norvegicus). Acta Neurobiol Exp 73, 430–7 (2013).

6. Behrens, T. E. J. et al. What Is a Cognitive Map? Organizing Knowledge for Flexible Behavior. Neuron 100, 490–509 (2018).

7. Brogden, W. J. Sensory pre-conditioning. J Exp Psychol 25, 323–332 (1939).

8. Lin, T.-C. E., Dumigan, N. M., Good, M. & Honey, R. C. Novel sensory preconditioning procedures identify a specific role for the hippocampus in pattern completion. Neurobiol Learn Mem 130, 142–8 (2016).

9. Talk, A. C., Gandhi, C. C. & Matzel, L. D. Hippocampal function during behaviorally silent associative learning: Dissociation of memory storage and expression. Hippocampus 12, 648–656 (2002).

10. Wong, F. S., Westbrook, R. F. & Holmes, N. M. “Online” integration of sensory and fear memories in the rat medial temporal lobe. Elife 8, e47085 (2019).

11. Holmes, N. M., Parkes, S. L., Killcross, A. S. & Westbrook, R. F. The Basolateral Amygdala Is Critical for Learning about Neutral Stimuli in the Presence of Danger, and the Perirhinal Cortex Is Critical in the Absence of Danger. J Neurosci 33, 13112–13125 (2013).

12. Todd, T. P., Huszár, R., DeAngeli, N. E. & Bucci, D. J. Higher-order conditioning and the retrosplenial cortex. Neurobiol Learn Mem 133, 257–264 (2016).

13. Sadacca, B. F. et al. Orbitofrontal neurons signal sensory associations underlying model-based inference in a sensory preconditioning task. Elife 7, 1–17 (2018).

14. Hart, E. E., Sharpe, M. J., Gardner, M. P. & Schoenbaum, G. Responding to preconditioned cues is devaluation sensitive and requires orbitofrontal cortex during cue-cue learning. Elife 9, e59998 (2020).

15. Robinson, S. et al. Chemogenetic Silencing of Neurons in Retrosplenial Cortex Disrupts Sensory Preconditioning. J Neurosci Official J Soc Neurosci 34, 10982–10988 (2014).

16. Fournier, D. I., Monasch, R. R., Bucci, D. J. & Todd, T. P. Retrosplenial cortex damage impairs unimodal sensory preconditioning. Behav Neurosci (2020) doi:10.1037/bne0000365.

17. Robinson, S., Keene, C. S., Iaccarino, H. F., Duan, D. & Bucci, D. J. Involvement of retrosplenial cortex in forming associations between multiple sensory stimuli. Behav Neurosci 125, 578–587 (2011).

18. Czajkowski, R. et al. Superficially projecting principal neurons in layer V of medial entorhinal cortex in the rat receive excitatory retrosplenial input. J Neurosci 33, 15779–15792 (2013).

19. Miller, A. M. P., Mau, W. & Smith, D. M. Retrosplenial Cortical Representations of Space and Future Goal Locations Develop with Learning. Curr Biol 29, 2083–2090 (2019).

20. Milczarek, M. M., Vann, S. D. & Sengpiel, F. Spatial Memory Engram in the Mouse Retrosplenial Cortex. Curr Biol 28, 1975–1980.e6 (2018).

21. Maguire, E. A. The retrosplenial contribution to human navigation: A review of lesion and neuroimaging findings. Scand J Psychol 42, 225–238 (2001).

22. Mitchell, A. S., Czajkowski, R., Zhang, N., Jeffery, K. & Nelson, A. J. D. Retrosplenial cortex and its role in spatial cognition. Brain Neurosci Adv 2, 2398212818757098 (2018).

23. Hattori, R., Danskin, B., Babic, Z., Mlynaryk, N. & Komiyama, T. Area-Specificity and Plasticity of History-Dependent Value Coding During Learning. Cell 177, 1858–1872 (2019).

24. Makino, H. & Komiyama, T. Learning enhances the relative impact of top-down processing in the visual cortex. Nat Neurosci 18, 1116–1122 (2015).

25. Auger, S. D. & Maguire, E. A. Retrosplenial cortex indexes stability beyond the spatial domain. J Neurosci Official J Soc Neurosci 38, 2602–17 (2018).

26. Chuong, A. S. et al. Noninvasive optical inhibition with a red-shifted microbial rhodopsin. Nat Neurosci 17, 1123–1129 (2014).

27. Akam, T. et al. The Anterior Cingulate Cortex Predicts Future States to Mediate Model-Based Action Selection. Neuron 109, 149–163.e7 (2021).

28. Blundell, P., Hall, G. & Killcross, S. Preserved Sensitivity to Outcome Value after Lesions of the Basolateral Amygdala. J Neurosci 23, 7702–7709 (2003).

29. Jacob, P. Y. et al. An independent, landmark-dominated head-direction signal in dysgranular retrosplenial cortex. Nat Neurosci 20, 173–175 (2017).

30. Voigts, J. & Harnett, M. T. Somatic and Dendritic Encoding of Spatial Variables in Retrosplenial Cortex Differs during 2D Navigation. Neuron 105, 237–245.e4 (2020).

31. Alexander, A. S. et al. Egocentric boundary vector tuning of the retrosplenial cortex. Sci Adv 6, eaaz2322 (2020).

32. Ennaceur, A., Neave, N. & Aggleton, J. P. Spontaneous object recognition and object location memory in rats: the effects of lesions in the cingulate cortices, the medial prefrontal cortex, the cingulum bundle and the fornix. Exp Brain Res 113, 509–519 (1997).

33. Vann, S. D. & Aggleton, J. P. Extensive Cytotoxic Lesions of the Rat Retrosplenial Cortex Reveal Consistent Deficits on Tasks That Tax Allocentric Spatial Memory. Behav Neurosci 116, 85–94 (2002).

34. Akam, T. et al. pyControl: Open source, Python based, hardware and software for controlling behavioural neuroscience experiments. Biorxiv 2021.02.22.432227 (2021) doi:10.1101/2021.02.22.432227.

